# Robust digital logic circuits in eukaryotic cells with CRISPR/dCas9 NOR gates

**DOI:** 10.1101/041871

**Authors:** Miles W. Gander, Justin D. Vrana, William E. Voje, James M. Carothers, Eric Kalvins

## Abstract

Natural genetic circuits enable cells to make sophisticated digital decisions. Building equally complex synthetic circuits in eukaryotes remains difficult, however, because commonly used genetic components leak transcriptionally, do not allow arbitrary interconnections, or do not have digital responses. Here, we designed a new *dCas9-Mxi1* based NOR gate architecture in *S. cerevisiae* that allows arbitrary connectivity and large genetic circuits. Because we used the strong chromatin remodeler *Mxi1*, our system showed very little leak and exhibits a highly digital response. In particular, we built a combinatorial library of NOR gates that each directly convert guide RNA (gRNA) input signals into gRNA output signals, enabling NOR gates to be “wired” together. We constructed and characterized logic circuits with up to seven independent gRNAs, including repression cascades with up to seven layers. Modeling predicted that the NOR gates have Hill Coefficients of approximately 1.71 ± 0.09, explaining the minimal signal degradation we observed in these deeply layered circuits. Our approach enables the construction of the largest, eukaryotic gene circuits to date and will form the basis for large, synthetic, decision making systems in living cells.

## Introduction

Living cells make decisions based on information processing genetic programs. Many of these programs execute digital functions^1–3^. The capability to build synthetic digital systems in living cells could allow engineers to build novel decision making regulatory networks for use in a variety of applications^4^, ranging from gene therapies that modify cell state based on sensed information, to entirely new developmental programs for crop species. In electronics a compositional approach has allowed the construction of digital circuits of great complexity to be quickly designed and implemented. Here, we have developed a framework that allows robust, digital circuits to be routinely constructed in eukaryotic cells.

Genetic components that implement simple logical operations, which in principle could be interconnected to form complex logic functions, have been demonstrated ^5–12^. DNA binding domains (DBDs) such as zinc fingers and TALEs have been used to construct libraries of transcription factors (TFs) in eukaryotes ^10,13,14^. However, scaling with DBDs has been difficult^15,16^. Recently, programmable and orthogonal CRISPR/dCas9 TFs^11,12,17–19^ have been employed to build up to five component circuits using CRISPR interference (CRISPRi) in prokaryotes^9^. Although CRISPRi allows programmable interconnections, CRISPRi is non-cooperative, leading to leak and signal degradation when layered^9^. Here, we address these issues, advancing the art of engineering living digital circuits by focusing on three main engineering goals.

First, we built a universal, single gene logic gate, in our case a NOR gate. It is well known that NOR gates can be composed to implement any logic function. Crucially, the input and output signals of our gates have the same molecular types while still being programmable so that, as in electronics, gates can be wired together. To achieve this, we made use of the CRISPR/dCas9 system: The signals in our framework are guide RNAs (gRNAs) whose sequences specifically match up to programmable target sequences on our NOR gate promoters. The NOR gate outputs are then gRNAs that match the target sequences on other NOR gate promoters. We avoided using RNA Pol III promoters to express gRNAs^11,17,18^ because they have low expression levels relative to Pol II promoters and are more difficult to engineer^20,21^. By programming the NOR gate input target sequences and output gRNA sequences in a set of gates, we were able to construct a variety of circuit topologies.

Second, we required that the digital “OFF” state for our gates corresponded essentially no expression of the output gRNA. To achieve this, we used the chromatin remodeling repression domain *Mxi1* to take advantage of the eukaryotic cell’s ability to repress gene expression, by fusing this domain to dCas9^22^. The *Mxi1* domain is thought to recruit histone deacetylase, and with it we observed strong transcriptional repression in our circuits^23^. Our results suggest that such repression provides a significantly improved “OFF” signal compared to CRISPR interference (CRISPRi), in which dCas9 is likely interfering with transcriptional initiation, but is not remodeling chromatin. The strong “OFF” behavior we observe with our NOR gates is a key factor that allows them to be composed into larger circuits by minimizing accumulation of transcriptional leak with every added layer.

Third, we required as digital a response as possible, which in this case means that as the concentration of the input gRNAs to a NOR gate go from low to high, the expression of the output gRNAs switches from high to low sharply, as opposed to gradually. Such a digital response is required so that NOR gates can be composed without the digital behavior of the resulting circuit degrading as the number of layers increases. Here, we employ the Hill coefficient, *n*, as a characterization of how digital our circuits are. From a mathematical model of our gates fit to both steady state and time response data, we show that that *n*≤1 results in degradation of the digital response with added layers of NOR gates. This behavior is what one would expect if we had used CRISPRi. On the other hand, when *n*>1 and even more so when *n* approaches 2, there exist parameters that allow our NOR gates to be composed without significant degradation. In particular, we estimate that *n*=1.71±0.09 (s.d.) with our NOR gates, and more importantly, show experimentally that we can build a variety of input logic circuits composed of up to five NOR gates and seven internal gRNA wires, as well as cascades of gates with up to seven layers that still have digital responses. The high Hill coefficient or cooperativity of our NOR gates is likely due to the complex recruitment of histone deacetylase by *Mxi1* and interactions with other proteins such as *SIN3*^24^ during repression. Thus, we did not specifically engineer the Hill coefficient in our system. Rather, we inherited it from our informed yet nevertheless fortuitous choice of the *Mxi1* repression domain.

In summary, we developed a framework in which single-gene NOR gates can be interconnected into arbitrary topologies that implement robust digital circuits in eukaryotic cells. Our approach allowed for the construction of the largest, eukaryotic gene circuits ever demonstrated. Because the technology is essentially generic and easy to rewire, it can in principle be used to implement arbitrary internal logic for a variety of synthetic cellular decision making systems, such as those being explored for diagnostics, therapeutics^25^, and development.

## NOR Gate Architecture

The gate NOR_i,j,k_, with input signals r_i_ and r_j_ and output r_k_, consists of a gRNA-responsive Pol II promoter (pGRRij) input stage, driving an output stage, ribozyme flanked gRNA (RGRk) (Fig. 1a). According to NOR logic, r_k_ is high only when both r_i_ and r_j_ are low. A signal, r_i_, is defined as a gRNA complexed with a *dCas9-Mix1* fusion protein, that confers strong transcriptional repression when bound to DNA^17^. The gRNA signals are distinguished by their unique 5′ guide sequence. A 20-component library of signals defining r_1_- r_20_ was used in this work (Extended Data 4). The pGRRij promoter contains two, 20 base-pair target sites that match r_i_ and r_j_ respectively. Since we designed twenty signals, there are 20^3^ = 8,000 total NOR gates in the set. A NOR_ij,k_ functions as a NOT_jk_ if the pGRR_ij_ contains two identical target sites, if the pGRR_i,j_ contains only one target site from the 20 component library (pGRR_i,null_), or if r_j_ is simply not used in the circuit. A target sequence of “null” refers to a pGRR that contains a target sequence that does not match any gRNA used in the containing circuit.

**Figure | 1.**
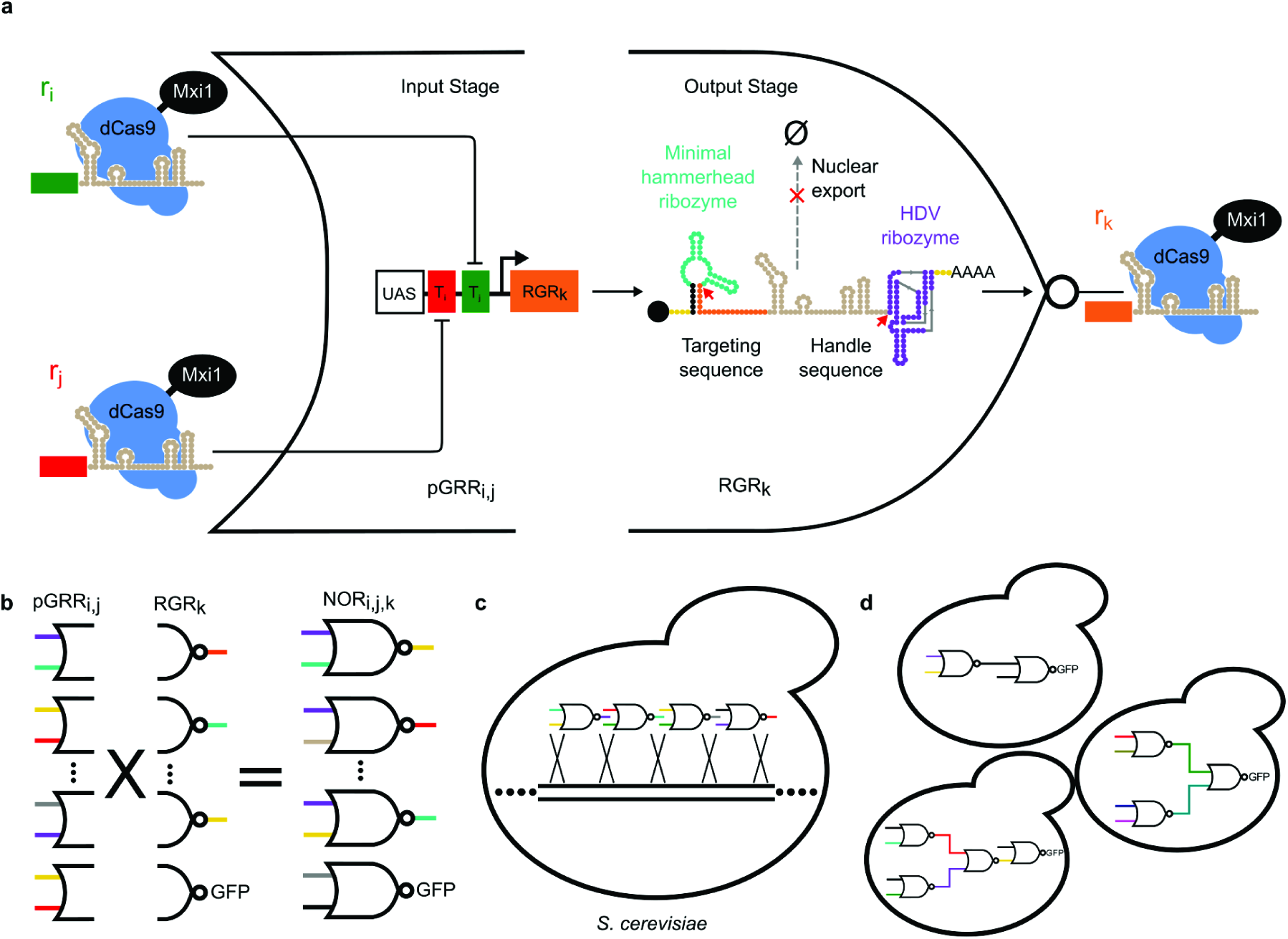
Schematic of the NOR gate architecture, library generation, and circuit composition. **a** A NOR gate comprised of an input stage consisting of a Pol II gRNA Responsive Promoter (pGRR) with two distinct gRNA target sites. The pGRR promoter is fully repressed by the binding of either one or both of its cognate gRNA:*dCas9-Mxi1* complexes. The output stage of the NOR gate is a gRNA transcript, flanked by self-cleaving ribozymes (RGR). Cleavage sites indicated by red arrows. The cleavage of the ribozymes prevents nuclear export of the gRNA, indicated by dotted grey arrow. **b** The process of NOR gate library construction. Our library consists of a set of 400 2-intput pGRR promoters and 20 RGR outputs, for a total of 8000 possible NOR gates. **c** Genomically integrating NOR gates into *S. cerevisiae*. **d** Arbitrary circuits are constructed by integrating multiple NOR gates into a single strain.

## Input Stage Promoter Design

The pGRR_i,j_ promoter is tightly repressed when gRNA:*dCas9-Mxi1* is bound to one or both of its two twenty base pair target sites. The core region of the pGRR_i,j_, the minimal pCYC1 promoter was chosen based on its successful use with dCas9 in the past^18^. Because the promoter has relatively low expression levels, and we wanted its output to be ON when not repressed, an upstream activating sequence (UAS), from the strong pGPD promoter^26^ was added, forming the base pGRR promoter. The UAS increased the unrepressed expression level of the pGRR output approximately three fold while maintaining the same OFF state expression level in the presence of r_i_ and r_j_, further separating the digital ON and digital OFF levels (Extended Data Fig. 5b). A library of 11 pGRR_i,j_ promoters, with i and j ranging from one to twenty, showed limited expression variability when driving *GFP*, with an ~ 18% standard deviation from the mean (Extended Data Fig. 5a). Sixteen of the twenty pGRR_i,null_:*GFP* constructs (i ranging from 1-20) were repressed to or near the level of *S. cerevisiae* autofluorescence in the presence of the corresponding signal r_i_ (Fig. 2b, Extended Data Fig. 3).

**Figure 2.**
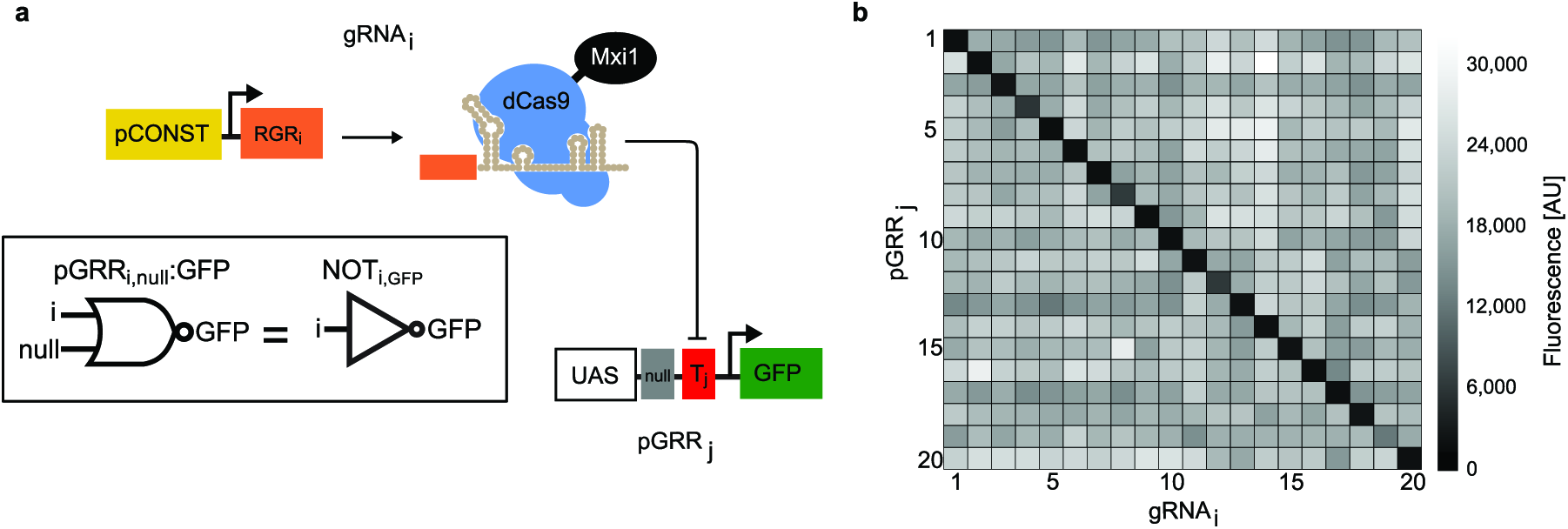
| RGR architecture and orthogonality of gRNA guide sequences. **a** Schematic of NOT gate used to test orthogonality of 20 gRNA and their cognate promoters. A constituitive promoter was used to express gRNA for these experiments. **b** Combinatorial library of gRNA and cognate promoters. Orthogonality of the computationally designed gRNA guide sequences was tested by crossing the 20 pGRR_i,null_ promoters, each expressing *GFP*, with the 20 gRNA_i_, creating 400 different strains of yeast. Fluorescence values of each strain were measured using flow cytometry. Fluorescence values from one biological replicate are displayed in the matrix. The heat map matrix shows strong repression with cognate pairs of pGRR_j_ and gRNA_i_ and minimal off-target repression.

**Figure 3.**
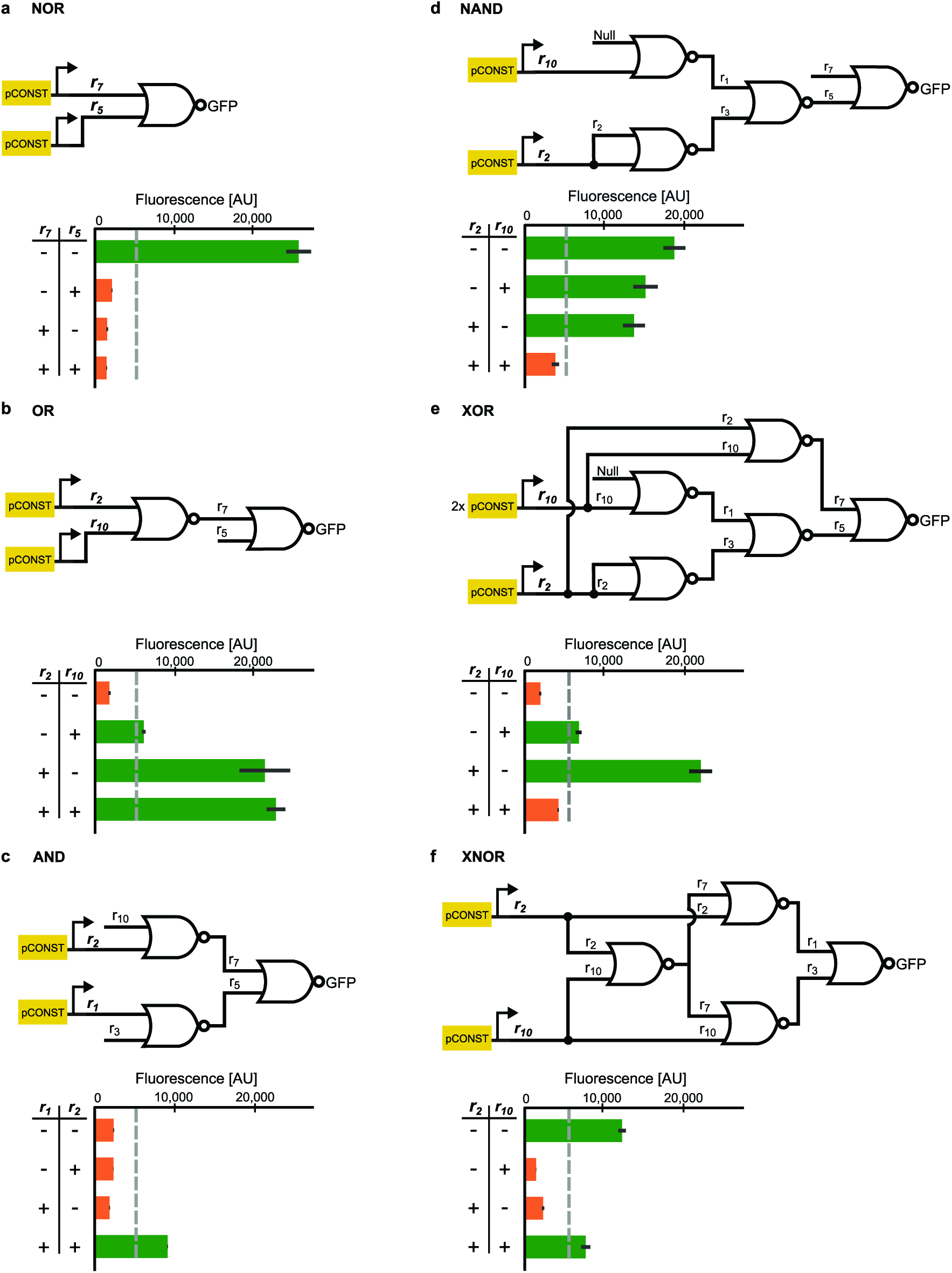
| NOR Gate-based logic circuits. **a-f** Six different two-input logic circuits constructed by interconnecting NOR gates. For each of the four input possibilities (- -, - +, + -, + +), a distinct strain was constructed with the corresponding inputs expressed off of constitutive promoters (for logical +), or not integrated at all (for logical -). Fluorescence values were collected using flow cytometry of cells growing in log phase. Error bars represent the standard deviation of three biological replicates measured during a single experiment.

**Figure 5.**
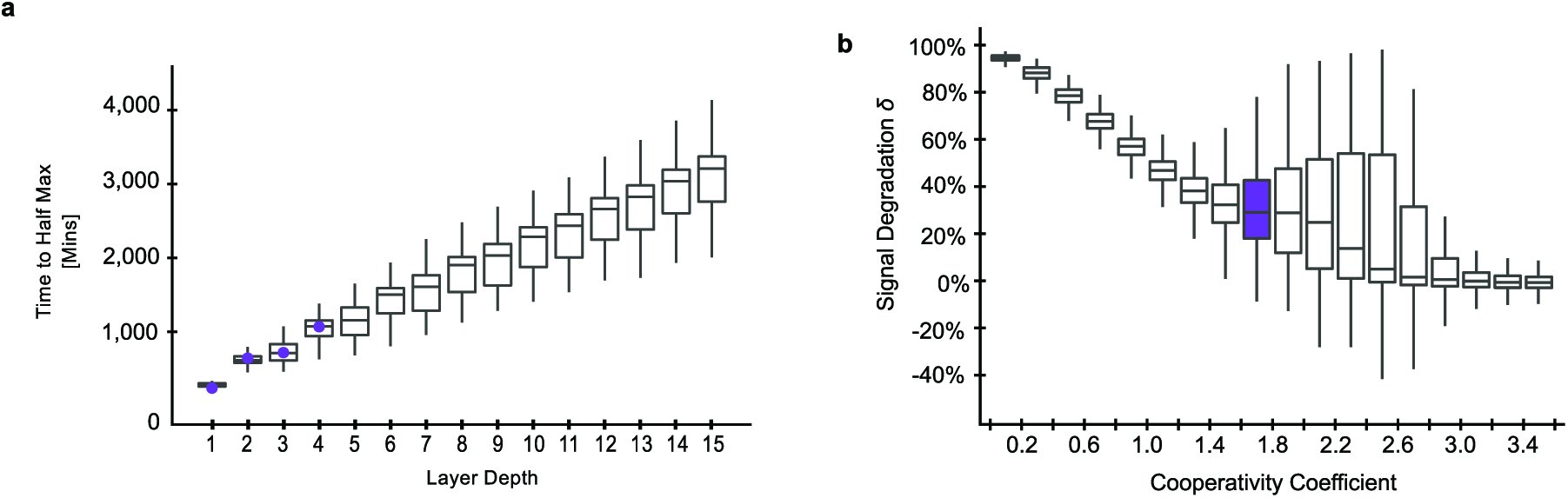
| Model predictions and analysis of repression cascades. **a** Simulations of time to half-maximal response using the model. Increasingly layered cascades show a positive linear relationship between circuit time to half-maximal response and circuit depth, with a slope of 197.67±0.45 (s.e.m.) mins/layer. The first four data points highlighted in purple are experimental data from Fig. 4b. **b** Signal degradation, *δ*, in a cascade decreases as cooperativity of *dcas9-Mxi1*increases. Boxplots of *δ* values were plotted with binned values of the cooperativity parameter *n*. As cooperativity increases, *δ*, on average decreases. At values of *n*> ~ 2.5, the median *δ* trends to zero. Between values of *n* =1.6 to *n* =2.8, the spread of *δ* is larger. The bin containing the predicted value of *dcas9-Mxi1, n* =1.71±0.09 (s.d.), is highlighted in purple.

## Output Stage RNA Design

Two different RNA pol II expression methods were used in this work (Extended Data Fig. 1). The first was an RGR design utilizing a 5′ minimal hammerhead ribozyme (mHH) and a 3′ Hepatitis delta virus ribozyme (HDV), flanking the gRNA^27^. The second was an “insulated” RGR (iRGR) with the mHH replaced by an avocado sunblotch viroid (ASBV) ribozyme. Both designs are intended to post-transcriptionally remove nuclear export signals, the 5′ cap and 3′ poly-A tail^28,29^. It has been shown that RNA device folding can be insulated from surrounding sequence context through computational sequence selection^30,31^. Both RGR architectures were computationally predicted to confer proper 5′ ribozyme folding for all 20 guide sequences. We observed similar levels of *dCas9-Mxi1* mediated repression with gRNAs expressed from both iRGR and RGR constructs (Extended Data Fig. 2). Interestingly, RGR transcripts lacking a 5′ ribozyme also showed *dCas9-Mxi1* mediated repression. These results are consistent with previous studies that indicate a majority of 5′ extended gRNA target sequences are processed to 20 nucleotides^32^. No significant crosstalk was observed when all r_1-10_(RGR design) and r_11-20_(iRGR design) were paired with all pGRR_1-20,null_:*GFP* among non cognate pairs (Fig. 2b, Extended Data Fig. 4). Sixteen out of twenty total RGRs (RGR_1-10_and iRGR_11-20_) when targeted to their cognate pGRR_1-20,null_:*GFP* constructs, repressed fluorescence to or near the level of autofluorescence for *S. cerevisiae* (Extended Data Fig. 3).

**Figure 4.**
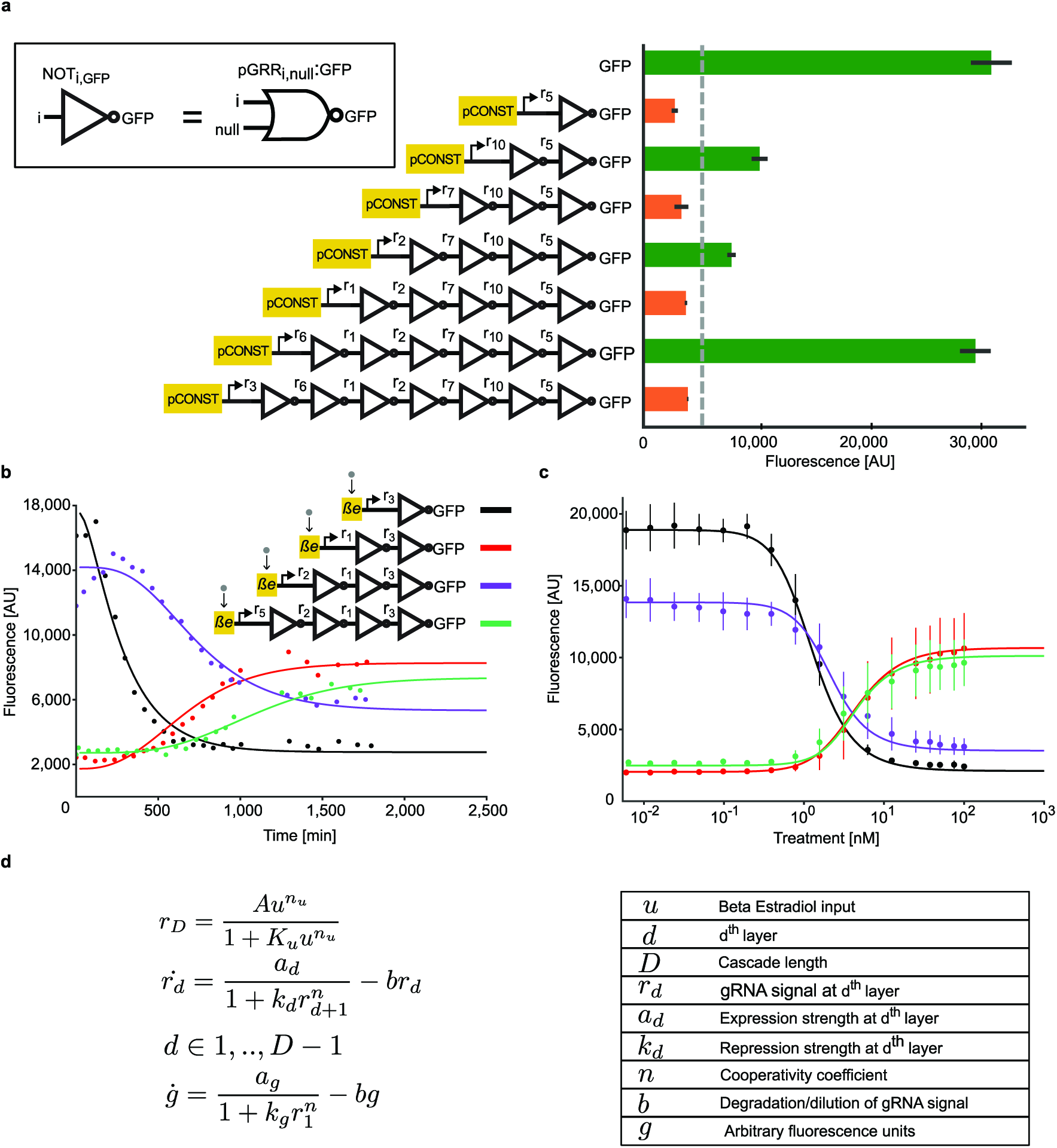
| Repression cascades characterization. **a**Repression cascades of one to seven gRNA. Cascades were created with sequential genomic integrations of NOT gates. The final output of each cascade is a NOT gate that expresses *GFP*. Each NOT gate represses the output of a subsequent NOT gate. Cascades with an even number of layers express a high level of *GFP*, creating a digital ON output, and odd depth cascades express low levels of *GFP*, creating a digital OFF output. Fluorescence measurements were taken using flow cytometry. Error bars represent the standard deviation of three biological replicates. **b**Temporal dynamics for cascades of one to four gRNA. Expression of the input gRNA was induced with Beta Estradiol. A model of the cascade, in which each layer is treated as a Hill function, was used to fit the data. The plot shows the data from one biological replicate. As the number of layers in the cascade increases, signal degradation and increased time to steady state is observed. **c**The steady state response function for the four inducible cascades. Error bars represent the standard deviation of three biological replicates measured over three separate experiments. **d**A representation of the model. The model was used to generate the fits for the steady state and kinetic inducible cascade experiments.

## Logic Circuits

Six two-input, one-output digital logic circuits were built by integrating up to five NOR gate cassettes into various selectable loci in the yeast genome (Fig. 3a-f). The output of each circuit was made observable by having the last NOR gate drive the expression of *GFP*. The circuits were constructed from the 16 guide sequences of the 20-component library that exhibited the strongest repression (Extended Data Fig. 3). The truth table for each gate was experimentally obtained by constructing four separate strains, one for each pair of possible input values, in which the corresponding gRNA input signals were expressed from constitutive promoters (Extended Date Table 2). To demonstrate the modularity and scalability of the NOR_i,j,k_ gates, the complex logic functions were built by extending simpler functions. For example, the AND circuit was built by wiring an inverter to the NOR gate driving *GFP* of the OR circuit. Although we observed fluorescence intensity differences in the digital ON and OFF states in various circuits, a single threshold value of 5,000 fluorescence units correctly distinguished the ON and OFF outputs for all circuits in a manner consistent with the corresponding truth tables.

## Cascades

Inverter cascades of depth one through seven were created with NOT gates (Fig. 4a). The cascade of depth *D* was made by the addition of a NOT gate to repress the input stage of the depth *D-* 1 cascade. Each successive addition of a NOT gate inverter resulted in switching the behavior of the output *GFP* expression, *g*. As seen previously with the two-input logic circuits, there is considerable variability within the ON and OFF states. However, circuits that are supposed to exhibit ON or OFF behavior are clearly distinguishable from one another when the threshold of 5,000 fluorescence units is applied. As cascade depth increased the fluorescence levels of the OFF states for all of the odd depth cascades increased. Similarly, except, for the cascade of depth 6, as cascade depth increased the fluorescence levels of the ON states for the even depth cascades decreased, suggesting a gradual degradation of circuit function as the number of layers increased.

To investigate the temporal characteristics of the inverter cascades, we analyzed the kinetics of cascades of depth one through four. The *β*-estradiol inducible promoter pGALZ4 was used to activate transcription of the input gRNA and *GFP* expression was periodically measured over the course of about 30 hours, while keeping the cells in log growth phase (Fig. 4b). With increasing cascade depth, a clear delay in output response was evident, with the cascades reaching half-maximal expression at 4.08±0.45, 10.78±1.04, 12.01±1.18 and 17.83±1.00 hours (ressd) for cascades of depth one through four respectively. The dose response curves of the four cascades were also measured after passaging cells over 5 days (Fig. 4c). Consistent with the steady state cascades, the induction of a gRNA targeting the input of the cascade switched the output of the cascade from OFF to ON (even depth cascades) or from ON to OFF (odd depth cascades). Some signal degradation with successive layers was observed (Fig. 4c), suggesting a limit to the possible depth of the cascades.

## Mathematical Modeling

A kinetic model was constructed to capture the behavior of our synthetic cascades. The model combines successive Hill functions to represent simple transcription and repression associated with each gRNA:*dCas9-Mxi1* signal. The parameters *α*_*d*_and *K*_*d*_roughly capture expression and repression strengths of the promoters driving each gRNA signal, *r*_*d*_. Parameters *n* and *b* capture the cooperativity of repression and degradation/dilution of gRN A: *dCas9-Mxi1* signals respectively (Fig. 4d). We assume that the parameter *n* is representative of the mechanism of *dCas9-Mxi1* repression so it was held constant across all gates in the model. The steady state dose response and kinetic inducible cascade data were both fit to the model (Fig. 4b-c). Due to the different growth conditions of the steady state and kinetic cascade experiments, two separate model fits were generated for each experiment. The cooperativity parameter, *n*, was determined using the steady state dose response data and was fixed for the kinetic data fit. The cooperativity *n* for the gRN A: *dCas9-Mxi1* was estimated from the dose response model fit to be *n* = 1.71±0.09 (s.d.).

The fitting results were found to correlate well with the experimental data. As predicted, the degradation/dilution term *b* increased in the kinetic data fit compared to the steady state dose response data (Extended Date Table 1). The rise in *b* represents a rise in dilution rate due to the log-phase growth rate during the kinetic cascade experiment compared to cell passaging in the steady state dose response experiment.

The temporal responses of the cascades were predicted from simulations using randomly sampled parameters within the range of the model fit. Parameter values for kinetic simulations were re-sampled from the model fit using the kinetic cascade experimental data. Response times were found to rise linearly with increasing circuit depth. Linear regression analysis showed each successive layer to increase the response time by an average of 197.67±0.45 (s.e.m.) minutes (Fig. 5a), consistent with our experimental results. Response delay was found to depend primarily on the degradation/dilution rate *b* of gRNA:*dCas9-Mxi1* (Extended Data Fig. 6), which controls the overall timescale of the dynamics.

To extrapolate the model to predict signal degradation for deeper cascades, cascades of 15 layers in depth with increasing values of *n* were analyzed via simulations using randomly sampled parameters within the range of the dose response fits. Parameter values for each layer were re-sampled from 144 different model fits. Dynamic range *ρd*, was calculated at each layer, *d*. Here dynamic range is defined for a cascade as the log fold-change of the maximal response of a cascade in response different levels of input gRNA, 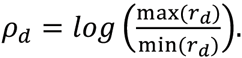A log-linear relationship was found between *ρ*_*d*_and *d*. This relationship was used to calculate the signal degradation, *δ*, representing the percent loss dynamic range per each additional layer.

Signal degradation was found to be largely dependent on the cooperativity coefficient, *n* (Fig. 5b). As cooperativity increases, *δ*, on average, decreases. At values of *n* > ~ 2.5, the median *δ* trends to zero. In the range of values of *n* =1.6 to *n=2.8* the spread of performance of the cascades is significantly larger. In this range the performance of the cascade is more sensitive to other parameters in the model. Our estimate of cooperativity, *n* =1.71±0.09 (s.d.), falls within the sensitive range, indicating the importance of utilizing well performing NOR gates in large circuits built using our architecture.

## Discussion

Our CRISPR/dCas9-based architecture represents a framework for constructing large, modular, transcriptional circuits. Other TF frameworks have suffered from lack of scalability and modularity, and signal degradation when layered. Our architecture addresses these issues by leveraging the orthogonality and modularity of the CRISPR/dCas9 system. To reduce signal leak and increase cooperativity, the strong transcriptional repression domain, *Mxi1*, was fused to *dCas9*. These basic advances resulted in some of the largest eukaryotic gene circuits, and the largest synthetic repression cascade, ever published.

While many circuit versions we constructed exhibited the correct truth table outcomes and appropriate switching behavior with addition of gRNA, some variants did not. We hypothesize that poor gate performance is due to two factors: the variance of expression between different pGRR_i,j_promoters and the inconsistency of the degree of repression of the pGRR_i,j_among different gRNA guide sequences (Extended Data Fig. 3, Extended Data Fig. 5). Indeed, the functional version of the XOR gate required double the expression of one of the inputs to exhibit correct XOR logic (Fig. 3e). Ultimately, the relationship between promoter sequence and expression level is an open question that requires further investigation^33^. Variation in repression level for different gRNA guide sequences could be due to either guide sequence effects on the flanking ribozyme cleavage rates, variations in DNA-RNA binding strength, or different association rates for the gRNA:*dCas9-Mxi1* complex^34,35^. Understanding the cause of these inconsistencies will allow for design of uniform components requiring fewer circuit variants to be screened before a properly functioning version is found.

Signal degradation effects were observed that would likely affect the desired function of larger synthetic circuits. Understanding and addressing the cause of the degradation is critical for more robust circuit design and construction in the future. One parameter of the system that is easily tunable is pGRR_i,j_promoter strength. Our model suggests that strong promoters will increase dynamic range of deeply layered circuits (Extended Data Fig. 6). This could be accomplished by adding additional UASs upstream of the pGRR_i,j_, or by reengineering pGRR_i,j_with nucleosome disfavoring sequences^21,36^.

According to our simulations increasing cooperativity, *n*, of the gRNA:*dCas9-Mxi1* will decrease the amount of signal degradation, *δ* (Fig. 5b). A key enabler of our system is the chromatin-remodeler Mxi1 that confers strong repression of target promoters. Chromatin remodeling TFs can achieve cooperativity through competition with nucleosomes^26^. This likely explains the increased cooperativity, *n* =1.71±0.09 (s.e.), of gRNA:*dCas9-Mxi1* as compared to similar systems using CRISPRi^9^. For values of cooperativity in the range *n* =1.6 to *n=2.8* the variance of *δ* is significantly larger than other cooperativity values. The increased variance indicates that signal degradation in the predicted cooperativity region is strongly correlated with NORijk performance. Low expression of gRNAk and incomplete repression of the pGRR_i,j_could confound circuit performance in the sensitive range. Our predicted *n* falls within the large variance region and could explain why some circuit variants using different NOR_i,j,k_, we constructed did not exhibit the correct truth tables. Thus, increasing cooperativity could allow for more robust circuit construction. Cooperativity could be increased in our architecture by engineering multi-domain protein interactions with *dCas9*, or via competition with decoy gRNA binding sites^37^.

The propagation time from input to output for deep circuits is an important consideration for understanding their applications. In electronics, for example, the propagation time determines the rate at which circuits can be “clocked”. Our model suggests that the propagation is strongly negatively correlated with the degradation/dilution rate *b*, of the *gRNA:dCas9-Mxi1* (Extended Data Fig. 6). For a simulated cascade with a depth of ten layers, the time to reach steady state is predicted to be on the order of two days (Fig. 5a). To decrease the time delay, degradation rate of gRNA:*dCas9-Mxi1* could be increased by adding ubiquitin tags to *dCas9* ^38^. However, our architecture may simply be more appropriate for complex slow-responding behaviors, like those found in cell-differentiation and development.

Biological systems utilize gene circuits to perform an abundant number of amazing functions, such as cell differentiation, metabolism and signal transduction. The ability to rationally engineer these functions, using synthetic digital circuits, will greatly impact many aspects of biotechnology. However, building such circuits with current technology is often exceedingly challenging. This work demonstrates that large eukaryotic circuits are indeed possible and is a step towards unlocking the vast potential of synthetic biology.

## Acknowledgements

Michelle Parks and the Klavins Lab technicians helped build strains and performed experimental assays. The computational work was facilitated though the use of the Hyak supercomputer system at the University of Washington. This research is funded in part by a grant from the Semiconductor Research Corporation and NSF Grant Number 1317653. J.M.C. was a fellow of the Alfred P. Sloan Foundation. W.E.V. and J.M.C. were supported in part by funds from the University of Washington and an NSF Award MCB 1517052.

## Author Contributions

M.W.G. and E.K. wrote the manuscript with contributions from all authors. M.W.G. and J.D.V. designed all experiments and performed data collection. J.D.V. developed and analyzed the mathematical model. W.E.V. performed model parameter sensitivity analysis. W.E.V. and J.M.C. performed all RNA design work.

## Author Information

The authors declare no competing financial interests. Correspondence and requests for materials should be addressed to M.W.G.

## Methods

### Construction of Yeast Strains

Yeast transformations were carried out using a standard lithium acetate protocol^1^into MATa W303-1A and MATalpha W303-1B. Matings of the MATa and MATalpha, were performed by coculturing both mating types and plating the culture onto selective agar media. All strains used in this work are detailed in Supplemental Table 2. All Sequences used for plasmid construction are available upon request.

### RNA design

RGR and iRGR sequences were computationally designed to enable the 5′ hammerhead ribozymes to fold into their target, functionally active, structures. ViennaRNA (RNAfold 2.1.9) was used to simulate long timescale (thermodynamic equilibrium) at an input temperature of 37C. Kinefold (kinefold_long_static_bianary 20060404) was used to simulate short timescale folding (co-transcriptional folding) with inputs of low and high polymerization rates of 25nt/s and 50 nt/s respectively, helix minimum free energy = 6.346 kcal/mol and folded without pseudoknots nor entanglements. 12 Kinefold simulations were run for each candidate sequence and agglomerated using custom python software to generate average folding trace data.

Ribozyme target structures needed for both viennaRNA and Kinefolds simulation evaluation were determined by folding ribozyme sequences (Minimal HH: 5′-NNNNNNCTGATGAGTCCGTGAGGACGAAACGAGTAAGCTCGTCNNNNNN −3′ ASBV1: 5′ - GGGACGGGCCATCATCTATCCCTGAAGAGACGAAGGCTTCGGCCAAGTCGAAACGGAAACGTCGGATAGTCGCCCGTCCC −3′) using RNAfold and Kinefold (melt and anneal of 1 minute), respectively. RGR targeting sequences and iRGR insulating sequences were screened in specific 5′ promoter contexts (pGAL1min: AGTATCAACAAAAAATTGTTAATATACCTCTATACTTTAACGTCAAGGAGAAAAAACTATACGGATTCTAGAACTAGTGGATCTACAAA, pAHD1: CAAGCTATACCAAGCATACAATCAACTATCTCATATACAGGATTCTAGAACTAGTGGATCTACAAA, pCYC1: ACTATACTTCTATAGACACACAAACACAAATACACACACTAATCTAGATATTGGATTCTAGAACTAGTGGATCTACAAA) and in the 3′ context of the targeting sequence and the gRNA handle sequence (gRNA handle: GTTTTAGAGCTAGAAATAGCAAGTTAAAATAAGGCTAGTCCGTTATCAACTTGAAAAAGTGGCACCGAGTCGGTGCTTTT).

Randomly generated 20 bp candidate targeting sequences for RGR, of which the most 5′ 6 bp defined the closing stem of the minimal HH ribozyme, were folded in the context of each promoter to confirm that the target structure was present in the MFE structure (viennaRNA) and that the target structure was present at >90% in the RNA folding trace at both low and high polymerase rates (Kinefold). Targeting sequences which enable correct folding in the context of each promoter were considered successful. For iRGRs, randomly generated 5′ and 3′ insulating sequences were designed for each of the three promoter types and were screened for function in the same manner. However, to select for the most robust insulating sequences each was screened against seventy-five randomly generated and ten randomly generated 20 bp guide sequences using viennaRNA and Kinefold, respectively.

### Cytometry

Fluorescence intensity was measured with a BD Accuri C6 flow cytometer equipped with a CSampler plate adapter using excitation wavelengths of 488 and 640 nm and an emission detection filter at 533 nm (FL1 channel). A total 10,000 events above a 400,000 FSC-H threshold (to exclude debris) were recorded for each sample with and core size of 22 mm using the Accuri C6 CFlow Sampler software. Cytometry data were exported as FCS 3.0 files and processed using the flowCore R software package and custom R scripts to obtain the mean FL1-A value at each data point. Scripts are available upon request.

### Data collection for orthogonality matrix

Cytometry readings were taken with cultures inoculated into synthetic complete with cells from freshly struck out on agar. Colonies were picked from plates and grown for 3 hours at 30° C before reads were taken.

### Data collection for logic circuits and static cascades

Cytometry measurements were taken on cells grown in cultures diluted 1:1000 from saturated culture, for 16 hours at 30°C.

### Data collection for inducible cascades

Cells from saturated culture were diluted 1:100 into fresh media with a Beta Estradiol (*βe*) concentration of 100nm. Cytometry measurements were taken over a ~ 30 hour period. During the time course, cells were periodically diluted to keep them in log growth phase. Experimental data was collected for steady state was measured for four strains, each containing four different *βe* inducible cascades. Each of the four strains was induced with eighteen different doses of *βe* ranging from 0 to 100 μM in a single batch of seventy-two cultures. Cells were diluted every eight to fifteen hours to prevent culture saturation. Steady-state fluorescence readings were taken after five days when the cultures were in log-phase.

### Model Description

A system of ODEs was used to model inducible repression cascades, with the dynamics of each gRNA modeled as Hill function dependent the concentration of another gRNA. The inducible gRNA (r_D_) was modeled using a beta-estradiol inducible promoter driving YFP, with YFP serving as a proxy for gRNA expression (data not shown). The equations below describe the model that was used in all downstream fitting procedures and analysis.

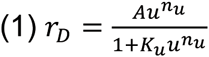

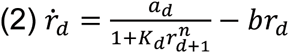

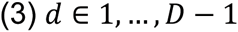

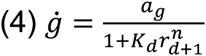

Parameters Descriptions:

*d*: *d* ^*th*^ layer

*D*: cascade length

*g*: arbitrary fluorescence (*AFU*)

*r*_*d*_: gRNA expression level at *d* ^*th*^ layer (*AU*)

*a*_*d*_: expression strength at *d* ^*th*^ layer (*AU* • *min* ^−1^)

*K*_*d*_: repressive strength at *d* ^*th*^ layer (*AU* ^−1^)

*b*: degradation/dilution of gRNA (*min* ^−1^)

*n*: co-operativity / Hill coefficient (unitless)

*u*: beta-estradiol input (*μM*)

*k*_*u*_: repressive strength at *d* ^*th*^ layer (*μM* ^−*n*_*u*_^)

*A*: expression strength of inducible gRNA (*AU* • *μM* ^−*n*_*u*_^)

### Fitting Procedure

Parameters were optimized using differential evolution (DE) followed by minimization using the Broyden-Fletcher-Goldfarb-Shanno (BFGS) algorithm^3^.

For the steady-state experiments, optimal parameter fits for the parameters *a*_0_^*ss*^ -*a*_3_^*ss*^, *k*_0_^*ss*^ -*k*_3_^*ss*^, *b* ^*ss*^, *u* ^*ss*^, and *n* were generated from three separate experiments. For each of the three experiments, 48 parameter fits were generated using DE/BFGS and means were calculated. The means from each experiment were used to determine the error (σ) for each parameter (Extended Date Table 1).

For the kinetics experiments, 38 parameter fits for *a*_0_^*kinetics*^ -*a*_3_^*kinetics*^, *k*_0_^*kinetics*^-*k*_3_^*kinetics*^, *b* ^*kinetics*^, *u* ^*kinetics*^ were generated from a single experiment. Means were calculated from this set of 38 (Extended Date Table 1). As there was only data for a single kinetics experiment, errors for the kinetic parameter values were unable to be calculated. The kinetics and steady-state parameter sets were resampled in downstream analyses to generate Monte-Carlo simulations of longer repression cascades.

## Extended Data

**Extended Data Figure 1.**
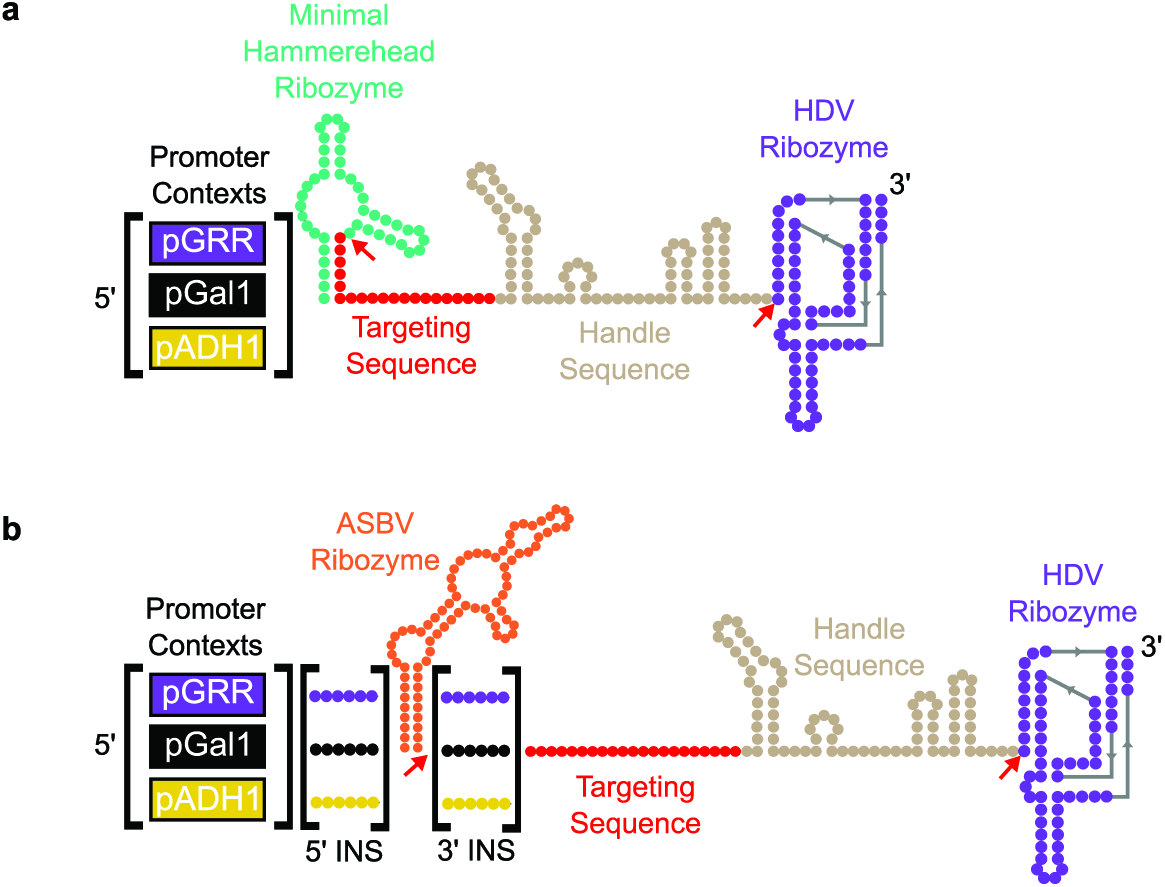
| Schematic of pol II gRNA expression systems. **a**The RGR architecture. All RGR constructs have targeting sequences that were computationally predicted to confer proper folding of the minimal hammerhead ribozyme in all three promoter sequence contexts used in the work. Cleavage sites are indicated by red arrows. **b**The insulated RGR (iRGR) architecture. The iRGR has unique 5′ and 3′ insulating sequences, designed for three promoter sequence contexts, flanking the ASBV ribozyme. In the presence of the insulating sequences, proper ASBV folding is predicted for the majority of targeting sequences. Cleavage sites are indicated by red arrows.

**Extended Data Figure 2.**
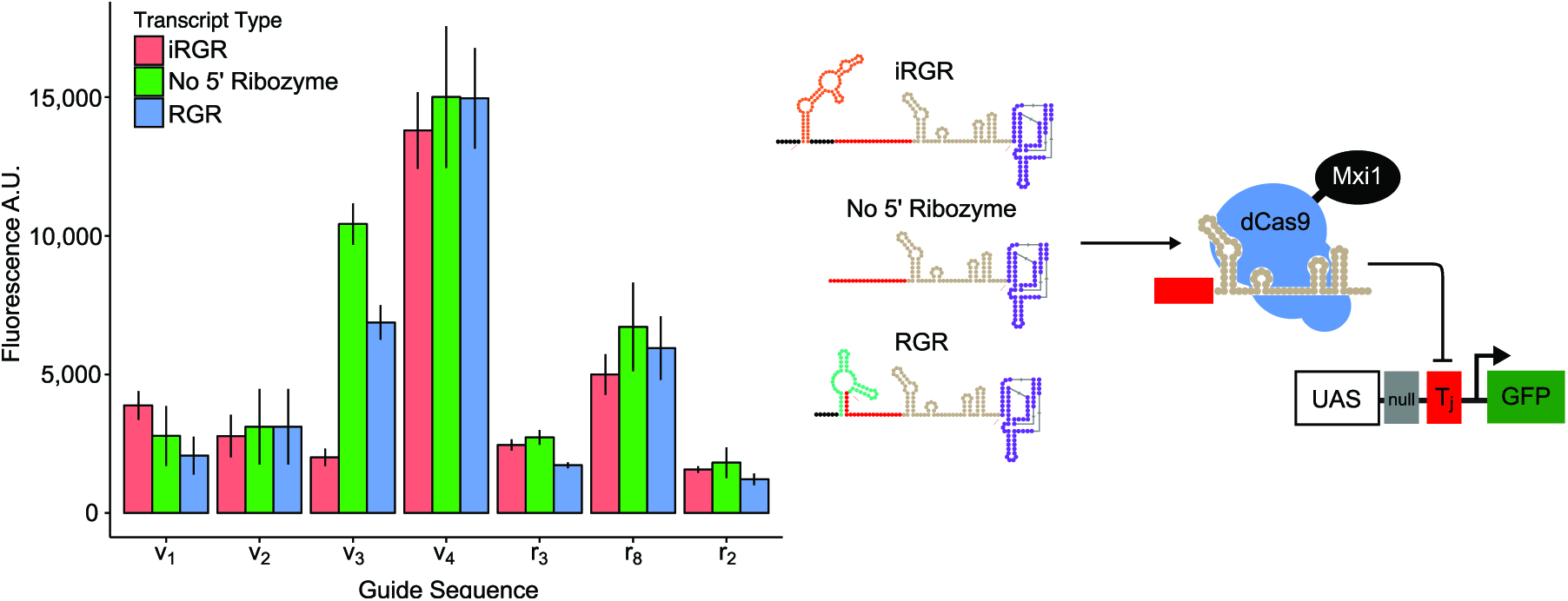
| Comparison of pol II gRNA expression designs. Seven gRNAs were expressed via three different designs, the RGR, the iRGR and an altered RGR design lacking the 5′ ribozyme. Guide sequences r_2_, r_3_, and r_8_were drawn from the gRNAs used in the main body of this paper, while guide sequences v_1_-v_4_were randomly generated guide sequences not contained within the original 20 component library. Fluorescence levels of repressed cognate pGRR promoters were measured via flow cytometry and error bars indicate standard deviation from 6 biological replicates, except for r_3_RGR, r_2_iRGR and r8 iRGR which represent 5 biological replicates. Data was collected across two different experimental runs. For all three transcript types, across all seven guide sequences except for v_3_, we observed comparable gRNA mediated repression of pGRR promoters. These data suggest that for many of guide sequences, the 5′ ribozyme is not a contributing factor in the behavior of the gRNAs in our system.

**Extended Data Figure 3.**
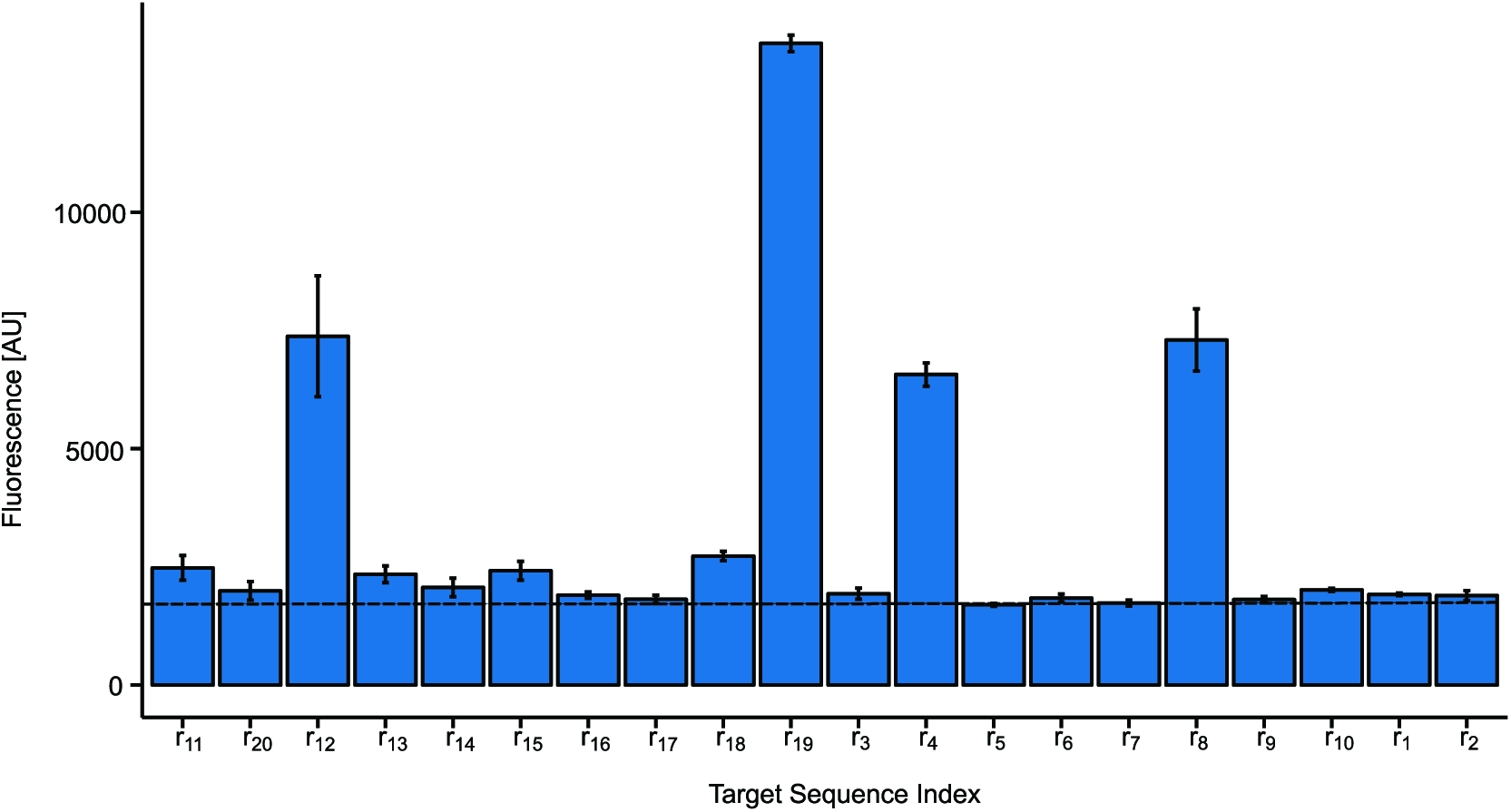
| Diagonal of orthogonality matrix repression variation. Bar chart representation of the diagonal of the orthogonality matrix from figure 2. Sixteen of the twenty guide sequences, when matched with their cognate promoter, show *GFP* repression near or at the level of autofluorescence for diploid *S. cerevisiae*. Autofluorescence, 1718.63 AU, indicated by the black dashed line. Four of the guide sequences exhibit significantly worse repression. The sixteen sequences that exhibit strong repression also exhibit variation in level of repression, indicating different levels of efficacy for each guide sequence. Possible causes for variable repression levels are discussed in the discussion. Error bars are standard deviation of fluorescence measurements from three biological replicates collected during one experimental run.

**Extended Data Figure 4.**
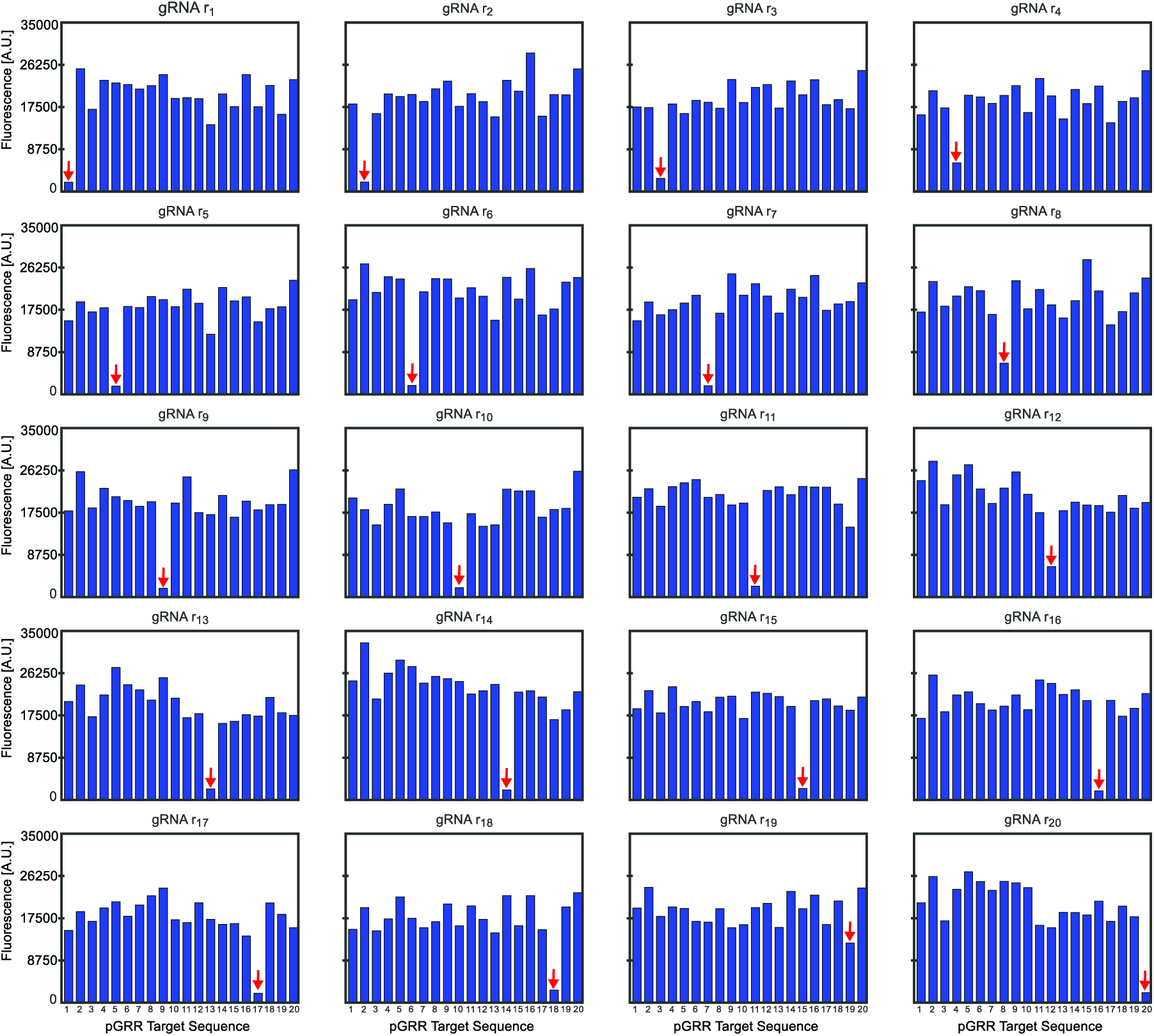
| Barchart of fluorescence values from orthogonality matrix. Flourescence values for all 400 strains in the orthogonality matrix. The strains are segmented by the 20 gRNA target sequences. Promoter target sequence index are in the same order for each subplot. Red arrows indicate a cognate pair of gRNA and pGRR promoter.

**Extended Data Figure 5.**
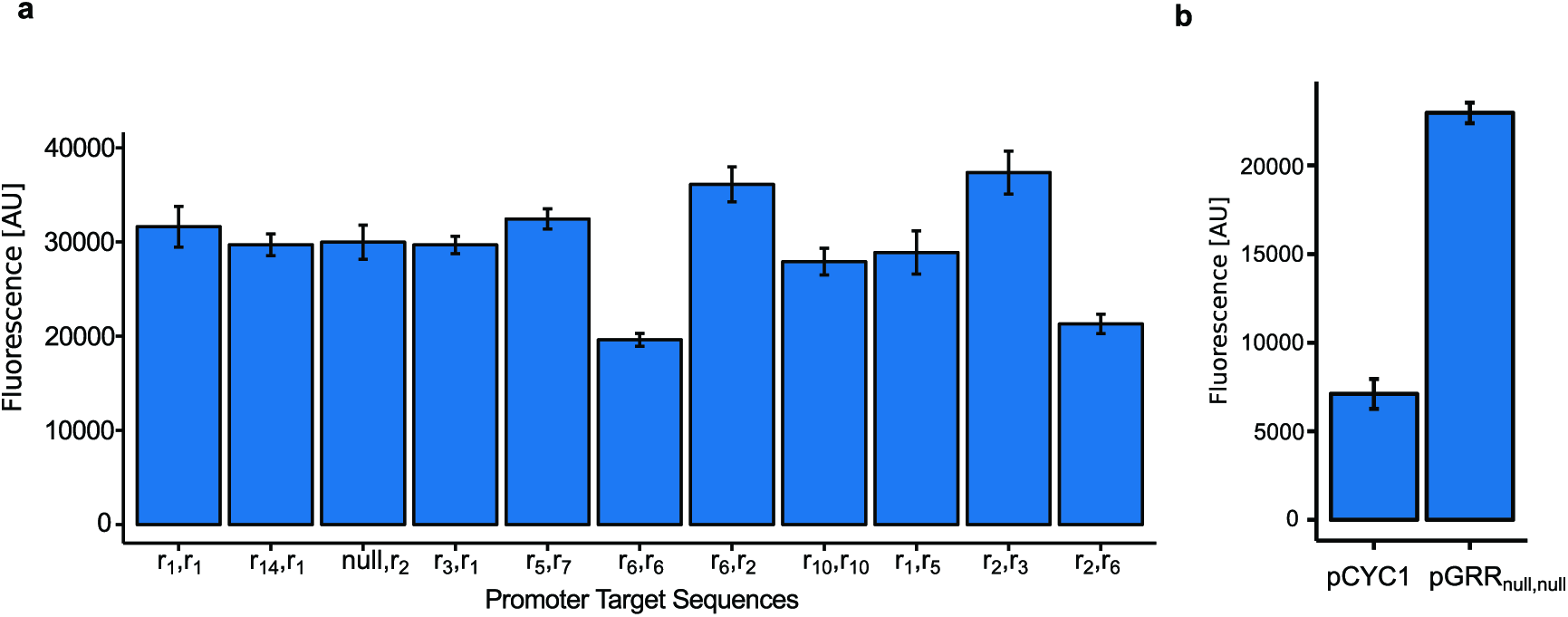
| pGRR promoter variability. **a**A subset of 11 pGRR_i,j_promoters driving *GFP* has a mean fluorescence of 29511.78 [AU], a standard deviation of 5357.249 [AU] and a range of 17751.67. Error bars represent the standard deviation of three biological replicates collected during one experimental run. **b**Addition of the pGPD UAS, to the pCYC1 minimal promoter, increases the expression of *GFP* 3.23 fold. Error bars represent the standard deviation of three biological replicates collected during one experimental run.

**Extended Data Figure 6.**
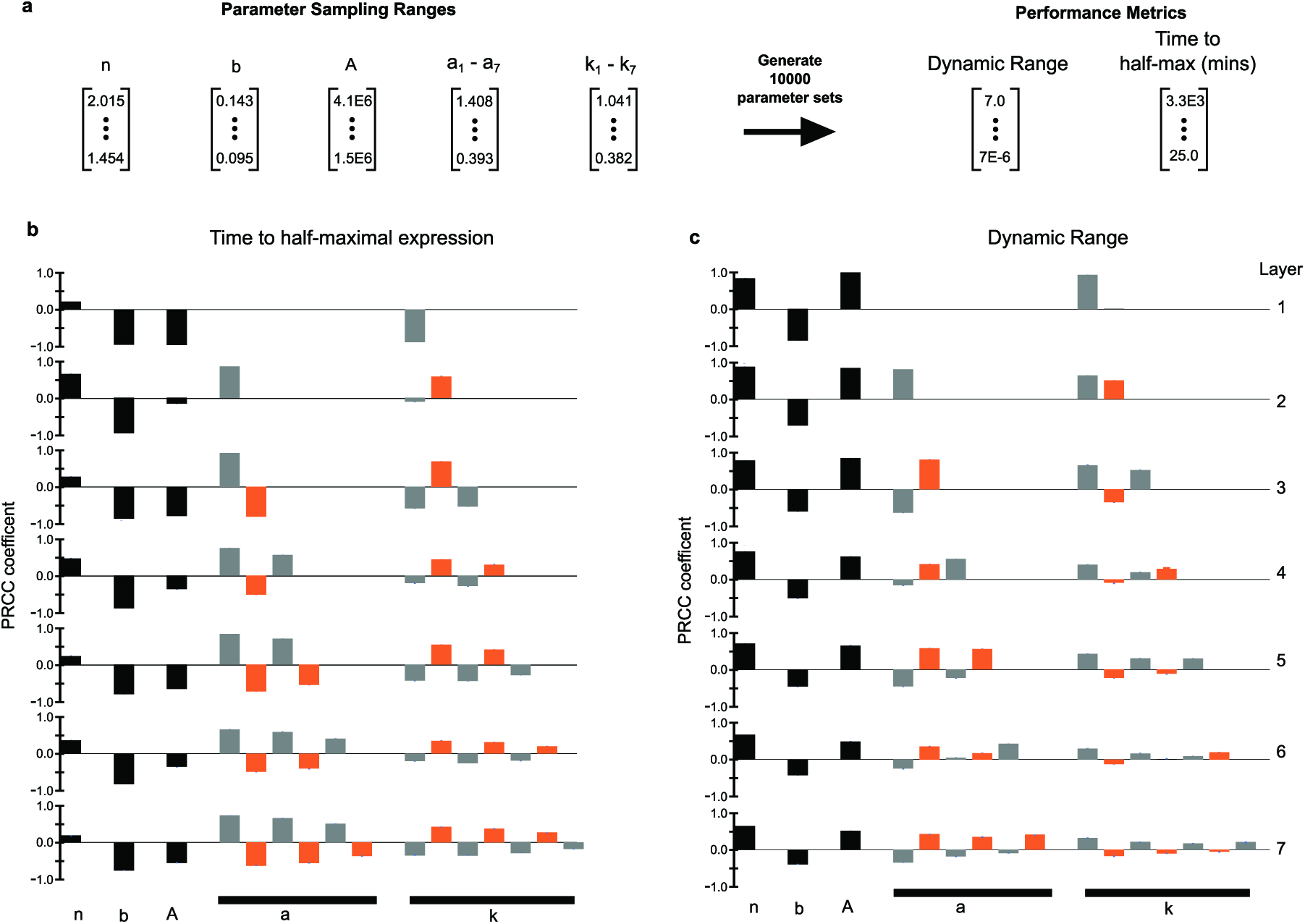
| Model Parameter Sensitivity. **a**10,000 parameter sets were resampled from a uniform distribution over the intervals shown and applied to our repression cascade model (see methods). **b**Partial rank correlation coefficient (PRCC) was used to determine the contribution of each parameter has on either dynamic range or the time-to-half max. PRCCs were calculated using R (R Foundation for Statistical Computing, Vienna, Austria) and 95% confidence intervals were estimated from bootstrap replicates (10 replicates with samples size 10,000). Parameters associated with odd and even layers are colored grey and orange respectively. At all layers in the time-to-half maximal plot, *b* very correlated with the output. In the dynamic range plot, *n*, strongly positively correlated at all layers with the output.

**Extended Data Table 1.** **| Parameter Fit Values**

| parameter | mean | std | units | description |
| --- | --- | --- | --- | --- |
| $a_g^{ss}$ | 1.408E+00 | 1.115E-02 | AU•min<sup>-1</sup> | expression strength of output promoter (steady state experiment) |
| $a_1^{ss}$ | 3.929E-01 | 7.370E-02 | AU•min<sup>-1</sup> | strength of promoter 1 (steady state experiment) |
| $a_2^{ss}$ | 5.278E-01 | 2.436E-02 | AU•min<sup>-1</sup> | strength of promoter 2 (steady state experiment) |
| $a_3^{ss}$ | 6.227E-01 | 1.436E-01 | AU•min<sup>-1</sup> | strength of promoter 3 (steady state experiment) |
| $k_g^{ss}$ | 1.041E+00 | 1.577E-01 | AU<sup>-n</sup> | repressibility of output promoter (steady state experiment) |
| $k_1^{ss}$ | 3.817E-01 | 1.669E-01 | AU<sup>-n</sup> | repressibility of promoter 1 (steady state experiment) |
| $k_2^{ss}$ | 4.006E-01 | 1.494E-01 | AU<sup>-n</sup> | repressibility of promoter 2 (steady state experiment) |
| $k_3^{ss}$ | 5.123E-01 | 1.328E-01 | AU<sup>-n</sup> | repressibility of promoter 3 (steady state experiment) |
| $A^{ss}$ | 2.824E+06 | 4.291E+05 | AU•uM<sup>-nu</sup> | induction strength of inducible promoter (steady state experiment) |
| $b^{ss}$ | 1.192E-01 | 7.913E-03 | min<sup>-1</sup> | gRNA degradation/dilution (steady state experiment) |
| $a_g^{kinetics}$ | 1.438E+00 | - | AU•min<sup>-1</sup> | expression strength of output promoter (kinetics experiment) |
| $a_1^{kinetics}$ | 4.265E-01 | - | AU•min<sup>-1</sup> | strength of promoter 1 (kinetics experiment) |
| $a_2^{kinetics}$ | 5.976E-01 | - | AU•min<sup>-1</sup> | strength of promoter 2 (kinetics experiment) |
| $a_3^{kinetics}$ | 5.504E-01 | - | AU•min<sup>-1</sup> | strength of promoter 3 (kinetics experiment) |
| $k_g^{kinetics}$ | 1.206E+00 | - | AU<sup>-n</sup> | repressibility of output promoter (kinetics experiment) |
| $k_1^{kinetics}$ | 5.285E-01 | - | AU<sup>-n</sup> | repressibility of promoter 1 (kinetics experiment) |
| $k_2^{kinetics}$ | 3.989E-01 | - | AU<sup>-n</sup> | repressibility of promoter 2 (kinetics experiment) |
| $k_3^{kinetics}$ | 2.930E-01 | - | AU<sup>-n</sup> | repressibility of promoter 3 (kinetics experiment) |
| $A^{kinetics}$ | 2.314E+06 | - | AU•uM<sup>-nu</sup> | induction strength of inducible promoter (kinetics experiment) |
| $b^{kinetics}$ | 1.306E-01 | - | min<sup>-1</sup> | gRNA degradation/dilution (kinetics experiment) |
| $n_u$ | 1.24E+00 | - | - | co-operativity of inducible promoter activation |
| $K$ | 7.318E+06 | - | uM<sup>-nu</sup> | inverse of dissociation constant |
| $n$ | 1.735E+00 | 9.361E-02 | - | co-operativity of dCas9-Mxi1 repression |

**Extended Data Table 2.** **| pCONST Promoter Table**

| Strain | pConst |
| --- | --- |
| Orthogonality Matrix Strains | pADH1:RGRi |
| Figure 3 NOR | pADH1:RGR-r7,pADH1:RGR-r5 |
| OR | pADH1:RGR-r2,pADH1 |
| AND | pADH1:RGR-r2,pGRR-r5:RGR-r1 |
| NAND | pADH1:RGR-r2,pGRR-nullnull:RGR-r10 |
| XOR | pADH1:RGR-r2,pADH1:RGR-r10,pGRR-nullnull:RGR-r10 |
| XNOR | pADH1:RGR-r2,pGRR-nullnull:RGR-r10 |
| StaticCascade 1 Layer | pGRR-r10:RGR-r5 |
| StaticCascade 2 Layer | pGRR-r7:RGR-r10 |
| StaticCascade 3 Layer | pGRR-r2:RGR-r7 |
| StaticCascade 4 Layer | pGRR-r1:RGR-r2 |
| StaticCascade 5 Layer | pGRR-r6:RGR-r1 |
| StaticCascade 6 Layer | pGRR-r3:RGR-r6 |
| StaticCascade 7 Layer | pGRR-r9:RGR-r3 |

**Extended Data Table 3.**
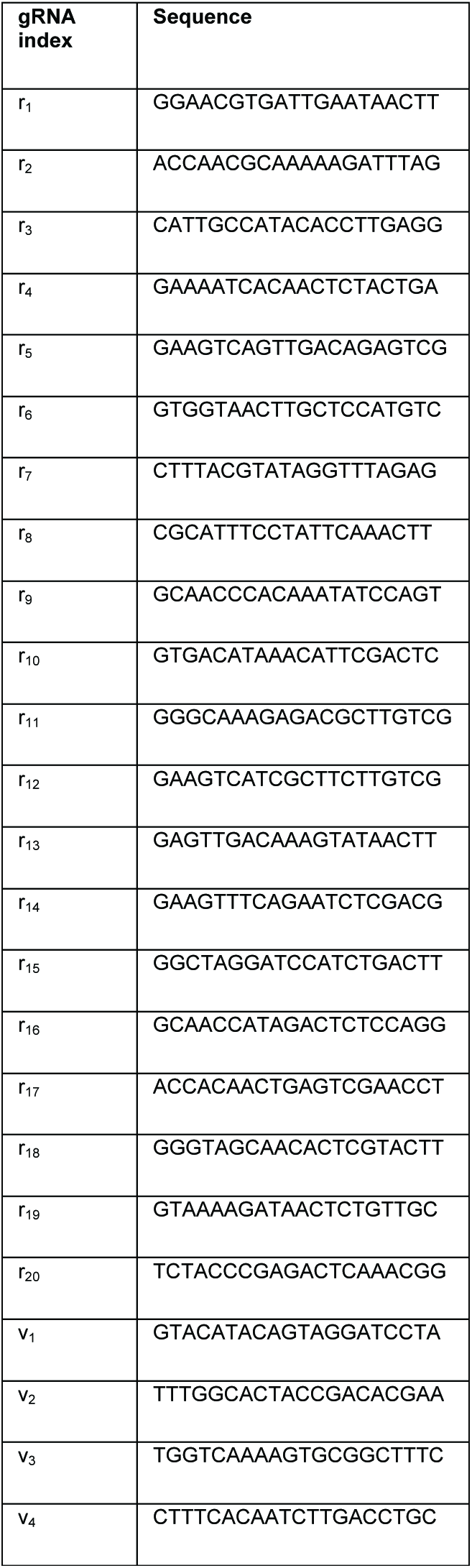
**| Guide Sequence Table**

## References

1. Shmulevich, Ilya, Edward R. Dougherty, and WeiZhang. “From Boolean to probabilistic Boolean networks as models of genetic regulatory networks.” Proceedings of the IEEE 90.11, 1778–1792(2002)

2. Watterson, Steven, StephenMarshall, and PeterGhazal. “Logic models of pathway biology.” Drug discovery today 13.9, 447–456 (2008).

3. Hinkelmann, Franziska, and ReinhardLaubenbacher. “Boolean models of bistable biological systems.” arXiv preprint arXiv:0912.2089 (2009).

4. Church, G. M., Elowitz, M. B., Smolke, C. D., Voigt, C. a & Weiss, R. Realizing the potential of synthetic biology. Nat. Rev. Mol. Cell Biol. 15, 289–94 (2014).

5. Lucks, J. B., Qi, L., Mutalik, V. K., Wang, D. & Arkin, A. P. Versatile RNA-sensing transcriptional regulators for engineering genetic networks. Proc. Natl. Acad. Sci. U. S. A. 108, 8617–8622 (2011).

6. Tamsir, A., Tabor, J. J. & Voigt, C. A. Robust multicellular computing using genetically encoded NOR gates and chemical ‘wires’. Nature 469, 212–215 (2011).

7. Moon, T. S., Lou, C., Tamsir, A., Stanton, B. C. & Voigt, C. A. Genetic programs constructed from layered logic gates in single cells. Nature 491, 249–253 (2012).

8. Siuti, P., Yazbek, J. & Lu, T. K. Synthetic circuits integrating logic and memory in living cells. Nat. Biotechnol. 31, 448–52 (2013).

9. Nielsen, A. A. K. & Voigt, C. A. Multi-input CRISPR / Cas genetic circuits that interface host regulatory networks. Mol. Syst. Biol. 1–11 (2014).

10. Stanton, B. C.et al.Genomic mining of prokaryotic repressors for orthogonal logic gates. Nat. Chem. Biol. 10, 99–105 (2014).

11. Kiani, S.et al.CRISPR transcriptional repression devices and layered circuits in mammalian cells. Nat. Methods 11, 723–6 (2014).

12. Didovyk, A. & Tsimring, L. Orthogonal Modular Gene Repression in Escherichia coli Using Engineered CRISPR/Cas9. (2016). doi:10.1021/acssynbio.5b00147

13. Klug, A. The discovery of zinc fingers and their development for practical applications in gene regulation and genome manipulation. Q. Rev. Biophys. 43, 1–21 (2010).

14. Zhang, F.et al.Efficient construction of sequence-specific TAL effectors for modulating mammalian transcription. Nat. Biotechnol. 29, 149–53 (2011).

15. Kittleson, J. T., Wu, G. C. & Anderson, J. C. Successes and failures in modular genetic engineering. Curr. Opin. Chem. Biol. 16, 329–336 (2012).

16. Joung, J. K. & Sander, J. D. TALENs: a widely applicable technology for targeted genome editing. Nat. Rev. Mol. Cell Biol. 14, 49–55 (2013).

17. Gilbert, L. a.et al.CRISPR-mediated modular RNA-guided regulation of transcription in eukaryotes. Cell 154, 442–451 (2013).

18. Farzadfard, F., Perli, S. D. & Lu, T. K. Tunable and multifunctional eukaryotic transcription factors based on CRISPR/Cas. ACS Synth. Biol. 2, 604–613 (2013).

19. Nissim, L., Perli, S. D., Fridkin, A., Perez-pinera, P. & Lu, T. K. Resource Multiplexed and Programmable Regulation of Gene Networks with an Integrated RNA and CRISPR / Cas Toolkit in Human Cells. Mol. Cell 54, 698–710 (2014).

20. Huang, Y. & Maraia, R. J. Comparison of the RNA polymerase III transcription machinery in Schizosaccharomyces pombe, Saccharomyces cerevisiae and human. Nucleic Acids Res. 29, 2675–2690 (2001).

21. Blazeck, J., Garg, R., Reed, B. & Alper, H. S. Controlling promoter strength and regulation in Saccharomyces cerevisiae using synthetic hybrid promoters. Biotechnol. Bioeng. 109, 2884–2895 (2012).

22. Schreiber-Agus, N.et al.An amino-terminal domain of Mxi1 mediates anti-Myc oncogenic activity and interacts with a homolog of the yeast transcriptional repressor SIN3. Cell 80, 777–86 (1995).

23. Lee, T. C. & Ziff, E. B. Mxi1 Is a Repressor of the c-myc Promoter and Reverses Activation by USF Mxi1 Is a Repressor of the c-myc Promoter and Reverses Activation by USF. J. Biol. Chem. 274, 595–606 (1999).

24. Mirny, L. A. Nucleosome-mediated cooperativity between transcription factors. Proc. Natl. Acad. Sci. 107, 22534–22539 (2010).

25. Xie, Zhen, et al.“Multi-input RNAi-based logic circuit for identification of specific cancer cells.” Science 333.6047(2011): 1307–1311.

26. Partow, Siavash, et al.“Characterization of different promoters for designing a new expression vector in Saccharomyces cerevisiae.” Yeast 27.11(2010): 955–964.

27. Gao, Yangbinand YundeZhao. “Self-processing of ribozyme flankded RNAs into guide RNAs in vitro and in vivo for CRISPR-mediated genome editing.” Journal of integrative plant biology 56.4, 343–349 (2014).

28. Lewis, J. D. & Izaurralde, E. The role of the cap structure in RNA processing and nuclear export. Eur. J. Biochem. 247, 461–469 (1997).

29. Dunn, E. F., Hammell, C. M., Hodge, C. A. & Cole, C. N. Yeast poly(A)-binding protein, Pab1, and PAN, a poly(A) nuclease complex recruited by Pab1, connect mRNA biogenesis to export. Genes Dev. 19, 90–103 (2005).

30. Carothers, J. M., Goler, J. a, Juminaga, D. & Keasling, J. D. Devices to Quantitatively Program. Science (80-.). 36498, 1716–1719 (2011).

31. Cambray, G.et al.Measurement and modeling of intrinsic transcription terminators. Nucleic Acids Res. 41, 5139–5148 (2013).

32. Ran, F. A.et al.Double nicking by RNA-guided CRISPR Cas9 for enhanced genome editing specificity. Cell 154, 1380–1389 (2013).

33. Sharon, E.et al.Inferring gene regulatory logic from high-throughput measurements of thousands of systematically designed promoters. Nat. Biotechnol. 30, 521–530 (2012).

34. Zoumadakis, M. & Tabler, M. Comparative analysis of cleavage rates after systematic permutation of the NUX consensus target motif for hammerhead ribozymes. Nucleic Acids Res. 23, 1192–6 (1995).

35. Fu, Y., Sander, J. D., Reyon, D., Cascio, V. M. & Joung, J. K. Improving CRISPR-Cas nuclease specificity using truncated guide RNAs. Nat. Biotechnol. advance on, (2014).

36. Curran, K. aet al.Design of synthetic yeast promoters via tuning of nucleosome architecture. Nat. Commun. 5, 4002(2014).

37. Brophy, J. A. N. & Voigt, C. A. Principles of genetic circuit design. Nat. Methods 11, 508–520 (2014).

38. Hochstrasser, M. Ubiquitin-Dependent Protein Degradation. Annu. Rev. Genet. 405–39 (1996).

## Methods References

1. Gietz, R. D. & Woods, R. a. Transformation of yeast by lithium acetate/single-stranded carrier DNA/polyethylene glycol method. Methods Enzymol. 350, 87–96 (2002).

2. Perez-Ortin, J. E., Alepuz, P. M. & Moreno, J. Genomics and gene transcription kinetics in yeast. Trends Genet. 23, 250–257 (2007).

3. Storn, R. & Price, K. Differential evolution-a simple and efficient heuristic for global optimization over continuous spaces. J. Glob. Optim. 11, 341–359 (1997).

